# Cannabinoid CB2 receptors enhance high-fat diet evoked peripheral neuroinflammation

**DOI:** 10.1101/2024.05.30.596629

**Authors:** Haruka Hosoki, Toru Asahi, Chihiro Nozaki

**Affiliations:** School of Advanced Science and Engineering, Waseda University, Tokyo, Japan; Comprehensive Research Organization, Waseda University, Tokyo, Japan; Research Organization for Nano & Life Innovation, Waseda University, Tokyo, Japan; Global Center for Science and Engineering, Waseda University, Tokyo, Japan

**Keywords:** Obesity, Neuroinflammation, Pain, CB2 receptors, endocannabinoid systems, CD11b+ monocytes

## Abstract

It is known that cannabinoid type 2 (CB2) receptor has anti-inflammatory role, therefore animals without CB2 receptors show enhanced inflammation and pain in the model of chronic pain e.g. neuropathic pain. We previously proposed the upregulated leptin signaling at the peripheral nerve as one of the underlying molecular mechanisms of pain exacerbation in nerve-injured CB2 knockouts, as they displayed robust upregulation of leptin receptors and leptin signaling in peripheral nerve. Due to these past results we hypothesized that CB2 receptor deficiency might also modify the peripheral neuroinflammation lead by chronic exposure to high fat diet (HFD). Interestingly, CB2 knockout animals showed the significant resistance to the HFD-induced neuroinflammation. Namely, 5-week feeding of HFD induced substantial hypersensitivity in WT animals, while tactile sensitivity of HFD-fed CB2 knockouts remained intact. HFD-fed WT animals also displayed the robust upregulation of chemokine CXCR4 expression with increased macrophage infiltration, which was never observed in HFD-fed CB2 knockout mice. Moreover, 5-week HFD-exposure lead significant increase of CD11b^+^Ly6G^-^Ly6C^high^ cells and decrease of CD11b^+^Ly6G^+^Ly6C^low^ cells in the spleen of WT animals, which was also not found in either HFD-fed CB2 knockouts or standard diet-fed WT and CB2 animals. Together with past report, these results suggest that CB2 receptors might have the double-sided regulatory role in context of the inflammation development, or more widely, immune system regulation. We propose that CB2 signaling is not always anti-inflammatory and could take pro-inflammatory role depending on the cause of the inflammation.

## Introduction

The interest in the medical use of cannabinoid ligands as the potential therapeutic agent for treatment of chronic pain and inflammation has greatly increased in the last decade[1]. Both cannabinoid receptor type 1 (CB1 receptors) and type 2 (CB2 receptors) are known to regulate pain in central and peripheral nervous system, however, how do they modify the pain is said to be different[2]. Thus, CB1 receptor-dependent analgesia is mediated by the presynaptic inhibition of GABAergic and glutamatergic neurons in supraspinal region[3], whereas CB2 receptors that is predominantly expressed in immune cells[4] work to reduce the pain by inhibiting the inflammatory response including peripheral neuroinflammation development. We previously investigated that animals lacking CB2 receptors displays enhanced neuroinflammation after the nerve injury, including development of mirror-image contralateral pain and dispersion of spinal activated microglias[5–7]. These results suggests that CB2 receptor activation may at least inhibit pain and neuroinflammation lead by peripheral nerve injury, and in fact several CB2 receptor agonists has been tested to show its analgesic effects in animal models of inflammation[8]. Interestingly, nerve-injured CB2 deficient animals showed robust upregulation of leptin receptor positive cells that lead enhanced leptin signaling at particular cells, which are the macrophages infiltrated in the nerve after the injury[6]. As blockage of leptin receptor signaling reversed the development of pain and neuroinflammation in CB2 knockouts [6], overall result suggested that one of the inflammatory pathways that CB2 receptor would calm down is the leptin receptor signaling, which highly contribute to the exacerbation of both neuroinflammation and neuropathic pain[9].

Because of these results, we then assumed whether deletion of CB2 receptor can modify the systemic inflammation lead by high-fat diet (HFD) induced obesity. It is well studied that chronic exposure to HFD evokes systemic inflammation including peripheral neuroinflammation and further hyperalgesia[10]. As HFD exposure also could modify the leptin signaling by increase of leptin release[11], we hypothesized that HFD-induced neuroinflammation might be enhanced by the deletion of CB2 receptors. However, we unexpectedly found that CB2 receptor might act rather pro-inflammatory during the chronic exposure of HFD; namely, HFD-fed CB2 receptor knockouts showed strong resistance to the peripheral neuroinflammation. Thus, 5-weeks exposure of HFD lead 1) tactile hypersensitivity, 2) significant macrophage infiltration and robust upregulation of CXCR4 in peripheral sciatic nerve, and 3) increase of CD11b^+^Ly6G^-^Ly6C^high^ cells and decrease of CD11b^+^Ly6G^+^Ly6C^low^ cells in the spleen, all of which were only observed in wild-type (WT) animals and never in CB2 knockouts. Interestingly, CB2 knockouts gained weight and blood glucose level by HFD as same as WT animals, suggesting that modulation of obesity-induced neuroinflammation by CB2 deficiency happened independently to the metabolism. Overall, present study shows that the role of CB2 receptor is not always to inhibit the inflammation as widely known, but may have pro-inflammatory function depending on the cause of the inflammation, such as HFD-induced obesity.

## Materials and Methods

### Animals

Different groups of male cannabinoid receptor 2 (CB2) knockout mice provided from Institute of Molecular Psychiatry, University of Bonn, Germany [12] with their wild-type (WT) littermate controls aging 6-weeks-old at the beginning of the experiments were used. CB2 knockout mice were backcrossed for more than 10 generations to C57BL/6J mice and were therefore congenic for this genetic background. Total of four cohorts for respective genotypes are prepared at the same time: Two cohorts for peripheral nerve injury group (sham and ligated) and two cohorts for HFD exposure group (HFD and standard diet, SD). Animals are collected from several breeding couples and two or more littermates are given for each cohort to avoid the litter effect. All mice were housed in a temperature (21 ± 1 °C) and humidity (55 ± 10%) controlled room with a 12-h light : 12-h dark cycle (light on between 08:00 h and 20:00 h). Food and water were available ad libitum except during behavioral observations. All experimental procedures and animal husbandry were carried out in accordance with the Law (No. 105) and Notification (No. 6) of the Japanese Government which is approved by the Committee for Animal Experimentation of the School of Science and Engineering at Waseda University (permission #A23-086) .

### Exposure to the high-fat diet (HFD)

Mice were housed in groups of two animals per cage. Body weight and blood glucose level were monitored once a week, over a 5-week high-fat or standard diet exposure period. The high-fat diet (HFD, D12492, OpenSource Diets, purchased from EP Trading, Tokyo, Japan) is consisted of 1.048 kcal/g protein, 3.144 kcal/g fat, and 1.048 kcal/g carbohydrates, with a total caloric value of 5.24 kcal/g. The standard diet (SD, D12450J, OpenSource Diets, purchased from EP Trading, Tokyo, Japan) is consisted of 0.77 kcal/g protein, 0.385 kcal/g fat, 2.695 kcal/g carbohydrates, with a total caloric value of 3.85 kcal/g. Food pellet was changed every 2-days to avoid oxidization of the food. Body weight was measured at 11:00 h and blood was collected from tail vein at the same time. Collected blood was then immediately used to measure the blood glucose level using portable glucose meter (Accu-Check Aviva, Roche Diagnostics GmbH).

### Induction of surgical neuropathic pain

Partial sciatic nerve ligation (pSNL)-induced neuropathic pain was produced by a tight ligation of approximately half the diameter of the left common sciatic nerve by 7-0 braid silk suture under deep inhalation anaesthesia (2.5% isoflurane) according to the surgical method described previously [13]. Experiments with sham mice followed the same surgical procedure except for the nerve ligation.

Note that sham surgery had no effect to the mechanical sensitivity assessed by following von-Frey method. Furthermore, pSNL animals as well as sham mice was housed in groups of two animals per cage with standard diet throughout entire experimental procedure.

### Assessment of mechanical sensitivity

The von-Frey filament up-down method [14] was used to evaluate the mechanical sensitivity. Briefly, mice were placed in a 8 cm x 10 cm x 15 cm Plexiglas box and the hindpaw plantar surface was gently probed with a series of eight von Frey filaments with logarithmically incremental stiffness (0.008, 0.04, 0.07, 0.16, 0.40, 0.60, 1.00 and 2.00 g, purchased from Bioseb, Vitrolles, France). Stimuli were presented by probing at intervals of 5-10 seconds, and sharp withdrawal or flinching of probed hindpaw was considered as a positive response. The threshold of the response (50% mechanical threshold) was calculated by using the up-down Excel program based on the equation formula described previously [14], which was generously provided by Allan Basbaum’s laboratory (UCSF, San Francisco, CA, USA). Mice were habituated to experimental environment for 3-days before each testing. Baseline responses were tested the day before the HFD exposure or the peripheral nerve injury. Tactile hypersensitivity was assessed once a week since the HFD exposure (HFD groups), or on 3, 7, 10 and 14 days postsurgery (pSNL groups).

### Immunohistochemistry

Following to the final behavioral experiment (End of 5-week HFD/SD exposure for HFD models or 14th postoperative day for pSNL Ligated/sham animals), four mice per group were deeply anesthetized with ketamine/xylazine (50/10 mg/kg, i.p.) and then intracardially perfused with ice-cold 0.1M phosphate buffer (PB), followed by 4% paraformaldehyde/PB. Both sciatic nerves (approx. 1 cm around the ligated area or the middle section of whole sciatic nerve) and dorsal root ganglia (DRG) from L4-L6 region were removed, postfixed in 4% paraformaldehyde for 2 h, and cryopreserved in 30 % sucrose/PB solution at 4°C. Tissues were then frozen in tissue-tek compound (Sakura Finetek) and cut to 10 μm thick longitudinal sections in a cryostat, mounted in Star frost-coated slides and kept under -80°C until further immunostaining.

The slides were incubated in anti-F4/80 antibody (1:100; Cedarlane) or anti-CXCR4 antibody (1:100; Abcam) at 4°C overnight, followed by incubation with Alexa488 or Alexa594 anti-goat secondary antibody (1:200; Invitrogen) and Alexa488 anti-rat secondary antibody (1:200; Invitrogen, for anti-F4/80) for 2 h at room temperature. After washing three times with PBS, the slides were mounted with FluoromountG containing DAPI. The sections were observed under the confocal microscope (LSM SP8, DMI 6000 CS, Leica). The images acquisition was done using LCS software (Leica) with a 63× objective. Quantification analysis of images are done using Fiji by ImageJ (NIH) to obtain %positive area to the entire tissue image.

### Flow Cytometry

Flow cytometry of splenocytes are used to confirm the modification of immune system by HFD as well as CB2 deletion. Single cell suspensions were prepared from freshly harvested spleen under isoflurane anesthesia using following procedure: First, spleens were mashed using a plunger and passed through a 70 μm filter with 0.5 mL 1xACK buffer. The cell strainer were washed twice with 0.5 mL 1xACK buffer, then incubated for 3 minutes in RT followed by 10 mL of ice-cold 0.5% BSA/PBS. Cell suspension were then centrifuged at 1500 rpm for 10 min, then the supernatant was discarded and cells are counted by Flow-count (Beckman Coulter). Nonspecific Fc binding of washed cells are blocked by with anti-mouse CD16/32 (2.4G2, BioXCell). Cell surface markers were then stained with the following antibodies: anti-mouse CD19-BV510 (GK1.5, Beckton Dickinson), CD5-BV421 (53-6.7, Beckton Dickinson), NK1.1-APC (PK136, Biolegend) to eliminate B cell, T cell and NK cell population, and CD11b-PerCP-Cy5.5 (M1/70, Beckton Dickinson) to pick up myeloid/monocyte population, then Ly6C-FITC (AL21, Beckton Dickinson) and Ly6G-eF450 (1A8, Thermo Fisher Scientific) to separate the subpopulation of CD11b+ myeloids/monocytes. Flow cytometry was performed on CytoFLEX S (Beckman Coulter), and the data (500000 events for spleen and bone marrow, 100000 events for peripheral blood) were acquired then analyzed using CytExpert software (Beckman Coulter).

### Statistics

The analysis of pharmacological/surgical or genotype effect was performed using three-way ANOVA (body weight, blood glucose level and time-course of behavioral tests) with Tukey’s multiple comparisons test or two-way ANOVA (summary of behavioral tests) with Sidak’s multiple comparisons test. The analysis of quantified images or flow cytometric data was performed using two-way ANOVA with Sidak’s multiple comparisons test. All statistical analysis was performed with Graphpad Prism (ver. 10.2.1).

## RESULTS

### CB2 receptor deficiency attenuates HFD-evoked peripheral hyperalgesia

It is known that high fat diet induces peripheral neuroinflammation and further hyperalgesia [10]. As such neuroinflammation development is strongly affected by the basal metabolism of animals, we first tracked the transition of weight gain (Fig 1A) and blood glucose level (Fig 1B) together with tactile sensitivity development (Fig 1C) during HFD exposure. As shown in the result, CB2 deficient animals gained the weight and blood glucose level equally as the WT controls throughout 5-week HFD feeding session (Fig 1A and B), suggesting that CB2 deficiency do not affect any major metabolic changes which could be evoked by 5-weeks HFD exposure. However, while WT animals developed significant tactile hyperalgesia by 5-week HFD feeding, CB2 knockouts did not show any hypersensitivity (Fig 1C and D), which was contradictory to our past findings on pain development in nerve-injured CB2 knockouts [5–7]. To confirm whether inflammatory response of CB2 deficient animals we used in current study are same as we have reported before [6], we further conducted the partial sciatic nerve ligation (pSNL) using the individual littermate animals which was housed with same diets as the control SD groups. As previously reported, those nerve-injured CB2 knockout animals showed significantly stronger hypersensitivity compared to WT controls, as well as mirror-image hyperalgesia on uninjured hindpaw (Fig 1E and F). Collectively, these results showed that CB2 deficient animals are sensitive to nerve-injury evoked pain but are rather resistant to HFD-induced hyperalgesia. These results also suggest that one of the molecular component for the development of HFD-evoked neuropathic pain might be activation of CB2 receptor signaling.

**Figure 1.**
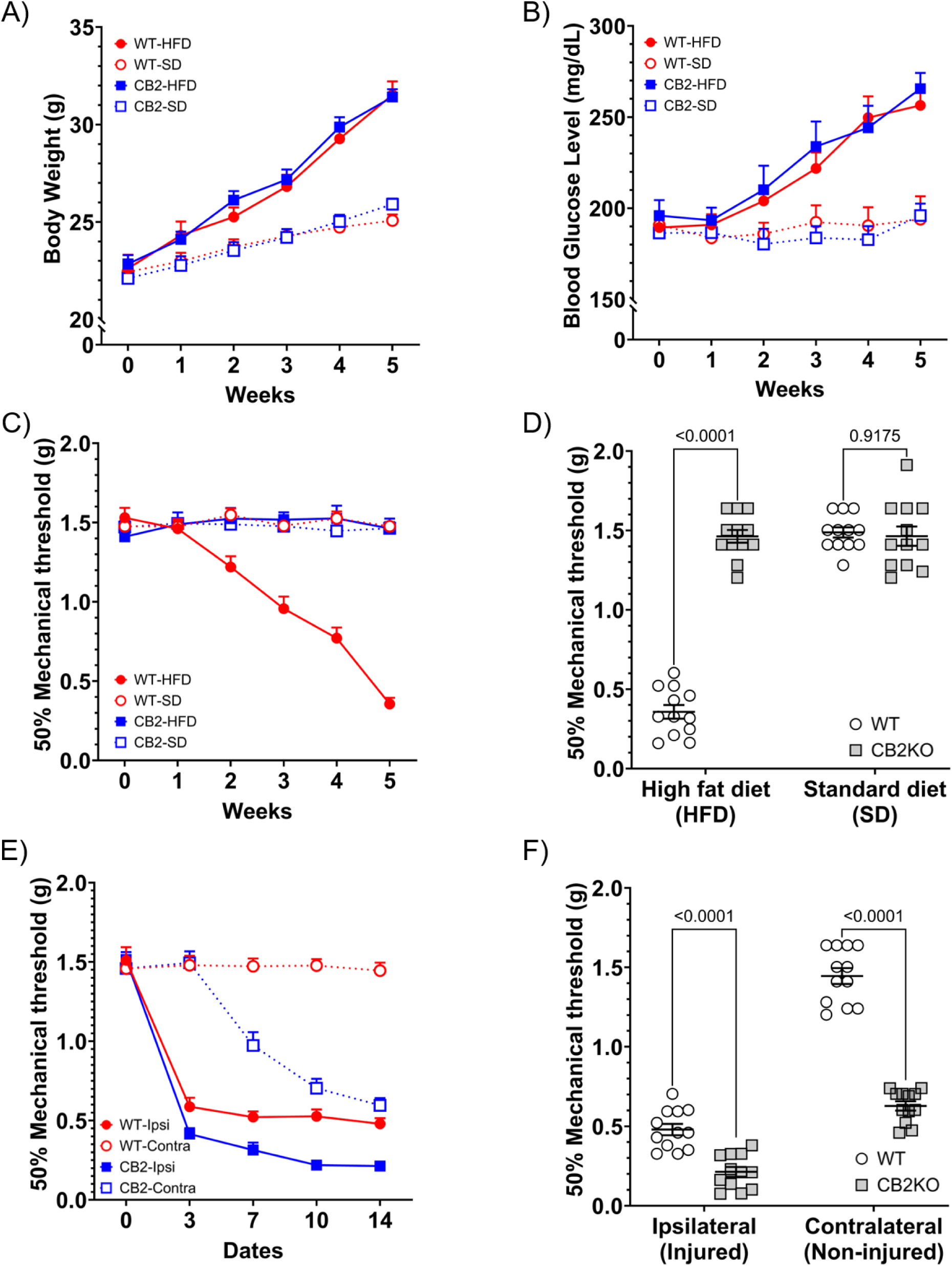
CB2 knockout mice are resistant to HFD-evoked hyperalgesia. While 5-weeks HFD exposure lead comparable gain of body weight and blood glucose levels in both WT and CB2 knockout animals, it only induced significant hyperalgesia in WT groups and not in CB2 knockouts. A, B) Both WT and CB2-knockouts showed similar weight and blood glucose level gain. C) WT developed hypersensitivity by 5-weeks HFD exposure, whereas HFD had no effect to CB2 knockouts. D) Pain threshold after 5-weeks HFD or SD exposure. Only HFD-fed WT showed significant hyperalgesia. E) CB2 deficiency enhanced nerve injury (ipsi)-induced hypersensitivity, together with development of mirror-image pain in uninjured contralateral (contra) paw. F) Pain threshold after 14-postoperative days in nerve-injured animals. CB2 knockouts showed enhanced pain development in both injured and uninjured paw. Significance is assessed by three-way ANOVA with Tukey’s multiple comparisons test (A-C and E, n=12) or two-way ANOVA with Sidak’s multiple comparisons test (D and F, n=12).

### CB2 deficiency inhibited HFD-evoked macrophage infiltration to peripheral nerves

Our previous reports showed that nerve-injured CB2 knockouts not only display stronger tactile hyperalgesia, but also the significant macrophage infiltration at peripheral nerve tissue [5, 6]. Here we conducted immunohistochemical analysis of F4/80 positive macrophages on sciatic nerve tissue in animal cohorts we used for the present study. We first could confirm that pSNL will certainly lead the significantly enhanced peripheral F4/80 positive macrophage infiltration at both injured and non-injured peripheral nerve tissue in CB2 deficient animals, compare to those of WT controls (Fig 2B). However, as seen in the previous pain experiment (Fig 1C and D), 5-week HFD-fed WT animals showed robust F4/80 positive macrophage infiltration in peripheral nerve, whereas HFD had almost no effect to the immunofluorescent staining in CB2 deficient animals (Fig 2A, p=0.9677 vs. SD-fed WT group). Note that standard diet also had no effect to peripheral neuroinflammation as well as the pain score measured before in both WT and CB2 knockouts.

**Figure 2.**
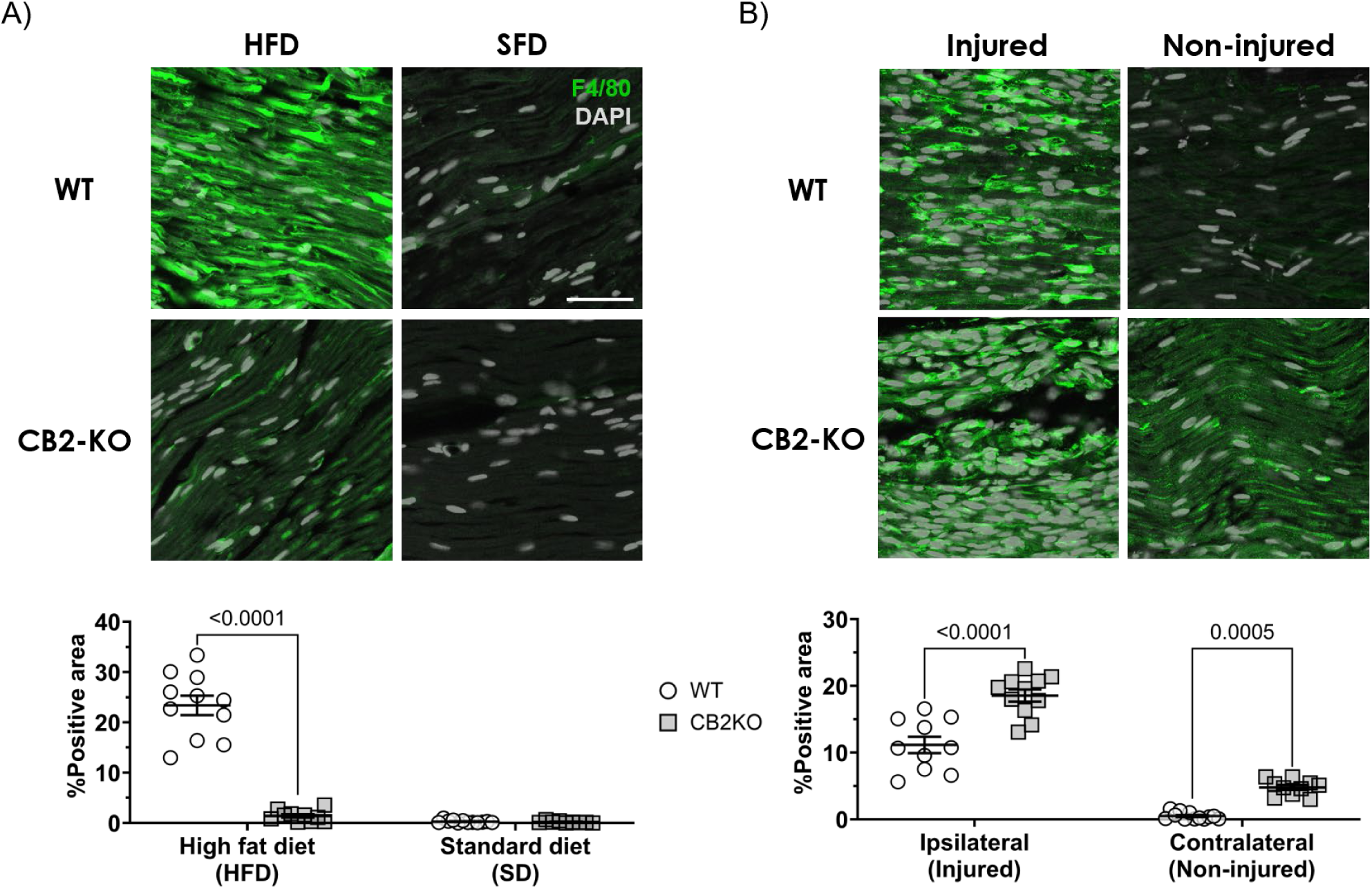
5-week HFD exposure lead peripheral macrophage infiltration in WT animals but not in CB2 knockouts. Macrophage infiltration to the peripheral nerve from nerve injured pSNL animals (A) and 5-week HFD exposed animals (B) has been evaluated by immunohistochemistry using F4/80 as the macrophage marker. Quantification of F4/80 positive cells (green) on each section are conducted by ImageJ software and quantified data are expressed as means ± SEM of 2-3 sections/4 mice/group. Bar on image indicates a length of 50 μm. Significant enhancement of macrophage infiltration has been observed in CB2 deficient pSNL animals (A) as well as WT animals after 5-week HFD exposure (B). Significance of quantified data was assessed by two-way ANOVA with Sidak’s multiple comparisons test (n=9-12).

Altogether, these results suggest that 5-week HFD feeding could evoke peripheral neuroinflammatory response only in WT animals which CB2 signaling is intact, but not in animals lacking CB2 receptors.

### CB2 deficiency enhances CXCR4 expression on the injured nerve, while it diminishes CXCR4 upregulation evoked by HFD

We further examined the contribution of other immunological factors such as chemokine receptors, and found that expression of CXCR4 is modified by CB2 deficiency. In nerve-injured neuropathic pain models, CB2 knockouts showed robustly enhanced CXCR4 expression compared to WT animals (Fig 3A). Such CB2 knockout-specific CXCR4 upregulation was observed not only at ipsilateral injured side, but also seen at non-injured contralateral side. Interestingly, upregulation of CXCR4 in WT nerve tissue was located at damaged region and almost no change was found at non-damaged region, whereas expansion of CXCR4 expression to such non-damaged region was observed in CB2 knockouts. On the other hand, HFD lead remarkable CXCR4 upregulation in peripheral nerve tissue of WT animals, but not in that of CB2 deficient mice (Fig 3B). Even after 5-weeks exposure of HFD, CB2 knockout nerve showed no significant CXCR4 upregulation compared to nerve tissue of SD-fed WT or CB2 deficient mice. These results indicate that while nerve injury-evoked CXCR4 upregulation can be inhibited by the presence of CB2 signaling, HFD requires CB2 receptor activity to enhance the CXCR4 expression in the peripheral nerve.

**Figure 3.**
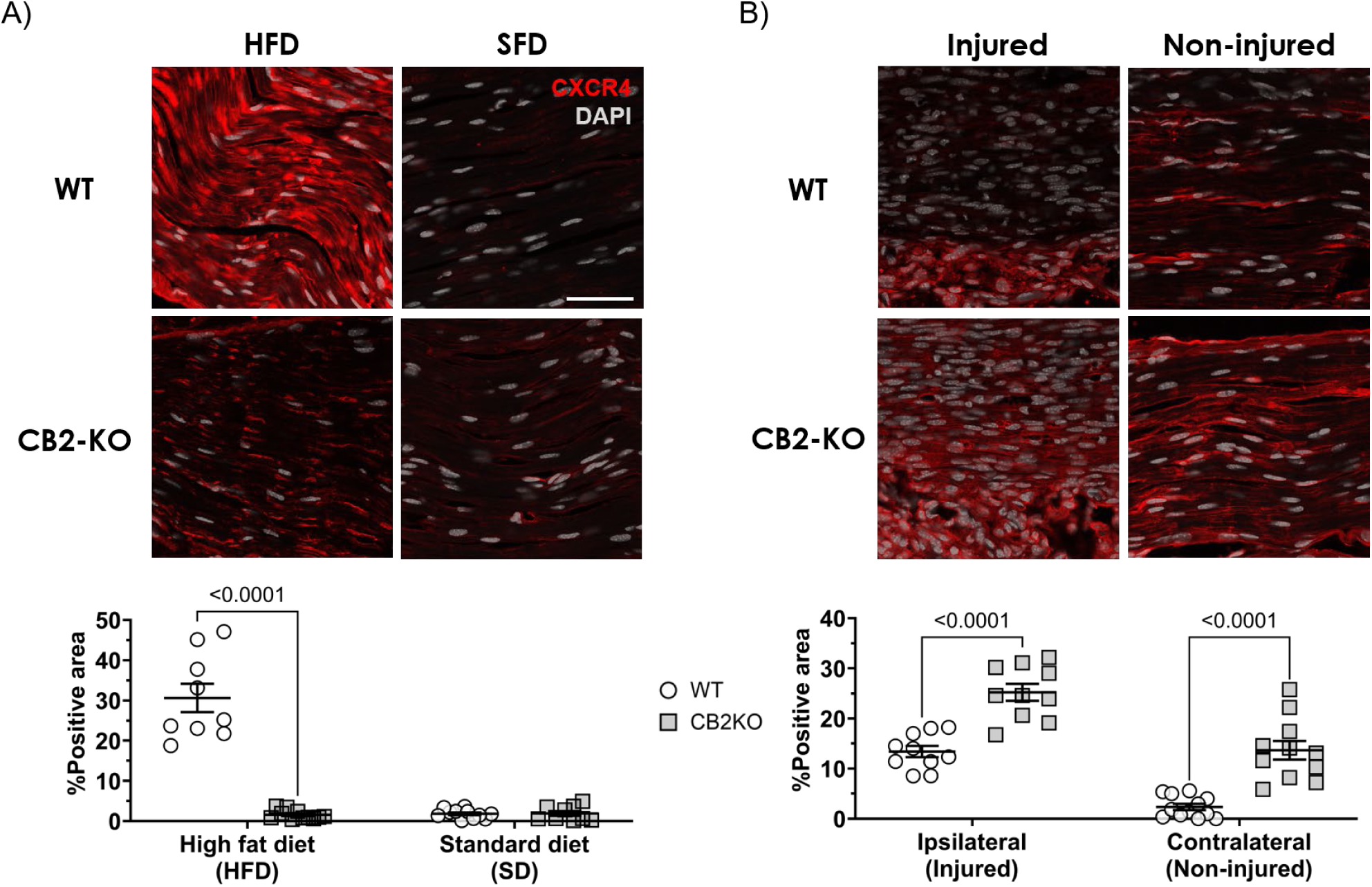
5-week HFD exposure lead significant CXCR4 upregulation in peripheral nerve tissue of WT animals but not in that of CB2 knockouts. CXCR4 expression of the peripheral nerve from nerve injured pSNL animals (A) and 5-week HFD exposed animals (B) has been evaluated by immunohistochemistry. Quantification of CXCR4 positive signals (red) on each section are conducted by ImageJ software and quantified data are expressed as means ± SEM of 2-3 sections/4 mice/group. Bar on image indicates a length of 50 μm. Significant enhancement of CXCR4 upregulation has been observed in CB2 deficient pSNL animals (A) as well as WT animals after 5-week HFD exposure (B). Significance of quantified data was assessed by two-way ANOVA with Sidak’s multiple comparisons test (n=9-12).

### CB2 deficiency also inhibited HFD-evoked CXCR4 expression on the dorsal root ganglia

Modification of CXCR4 expression by lack of CB2 activity has also been found in afferent region of peripheral nerve, which is dorsal root ganglia (DRG). As observed in sciatic nerve tissue, significant upregulation of CXCR4 positive signal evoked by nerve injury has been seen in CB2 deficient DRG tissue (Fig 4A). On the other hand, 5-week HFD exposure lead enhanced CXCR4 expression in WT DRG, while CB2 knockout showed only weak CXCR4 upregulation compare to SD-fed groups (Fig 4B). Thus, CXCR4 level in DRG will increase in CB2 knockout under the nerve injury, whereas it increase in WT groups under the 5-week HFD exposure. These results indicates that significant CXCR4 upregulation which was observed in peripheral nerve corresponds to them of DRG.

**Figure 4.**
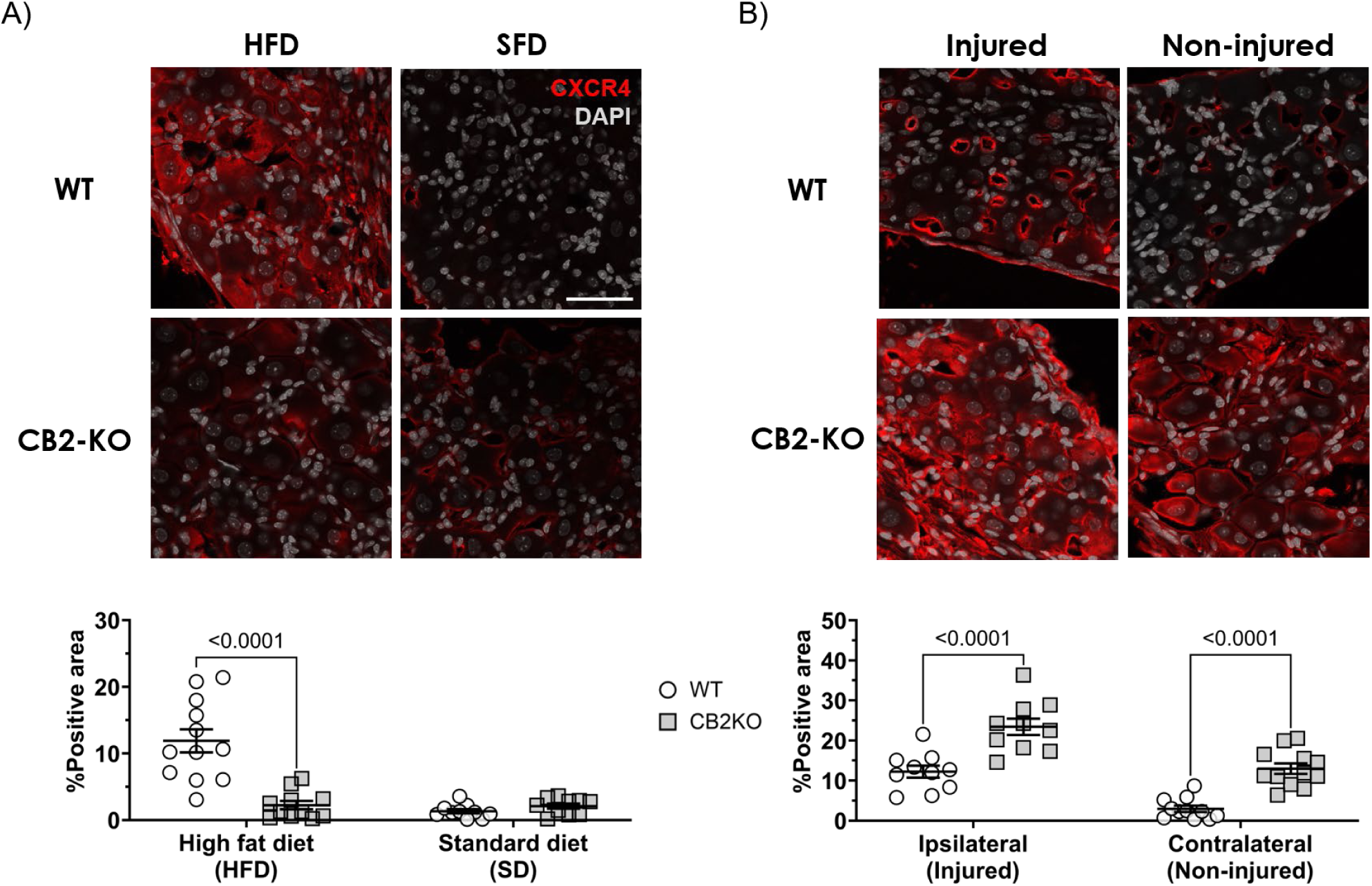
5-week HFD exposure lead significant CXCR4 upregulation in DRG of WT animals but not in that of CB2 knockouts. CXCR4 expression of DRG from nerve injured pSNL animals (A) and 5-week HFD exposed animals (B) has been evaluated by immunohistochemistry. Quantification of CXCR4 positive signals (red) on each section are conducted by ImageJ software, and quantified data are expressed as means ± SEM of 2-3 sections/4 mice/group. Bar on image indicates a length of 50 μm. Significant enhancement of CXCR4 upregulation has been observed in CB2 deficient pSNL animals (A) as well as 5-week HFD-fed WT animals (B). Significance of quantified data was assessed by two-way ANOVA with Sidak’s multiple comparisons test (n=10-12).

### HFD-feeding modulates the amount of splenic CD11b^+^ myeloid subpopulation in WT animals but not in CB2 deficient mice

Immunostaining of peripheral nerve and DRG showed the enhancement of HFD-lead neuroinflammation in WT can be due to excess infiltration of macrophages together with upregulation of CXCR4 chemokine receptors, however how HFD can evoke such inflammation only in WT and not in CB2 knockouts was unclear. Also which immune cells out of macrophage might contribute to such CB2-mediated inflammatory response is not clarified as well. Flow cytometry has been therefore conducted to investigate whether there are immune cell populations that could be modulated specifically in HFD-fed WT mice and not in HFD-fed CB2 animals (Fig5 and 6). We first found that HFD exposure had no effect to the total number of splenic T cells (CD5^+^), B cells (CD19^+^) and NK cells (NK1.1^+^) regardless of genotype. Furthermore, genotype difference also had no effect to total amount of splenic myeloid cells (CD5^-^CD19^-^NK1.1^-^CD11b^+^, Fig 5A). However, subpopulation of splenic myeloid cells has been largely modified by HFD in WT animals (Fig 5B and C). Thus, myeloid subset which is Ly6C^low^Ly6G^+^ cells has been decreased while another subset Ly6C^high^Ly6G^-^ cells has been increased in HFD-fed WT animals. Strikingly, neither decrease of Ly6C^low^Ly6G^+^ subsets nor increase of Ly6C^high^Ly6G^-^ subsets happened in HFD-fed CB2 knockout group. SD exposure showed no genotype effect, considering that the ratio of basal CD11b^+^ myeloid subsets was almost the same in WT and CB2 knockouts. Notably, distribution pattern of myeloid subpopulation in HFD-fed CB2 knockouts were comparable to that of either SD-fed CB2 knockouts or WT animals. These results indicate that lack of CB2 receptors might inhibit the irregular differentiation of splenic CD11b^+^ myeloid subpopulation which is evoked by 5-week HFD feeding.

**Figure 5.**
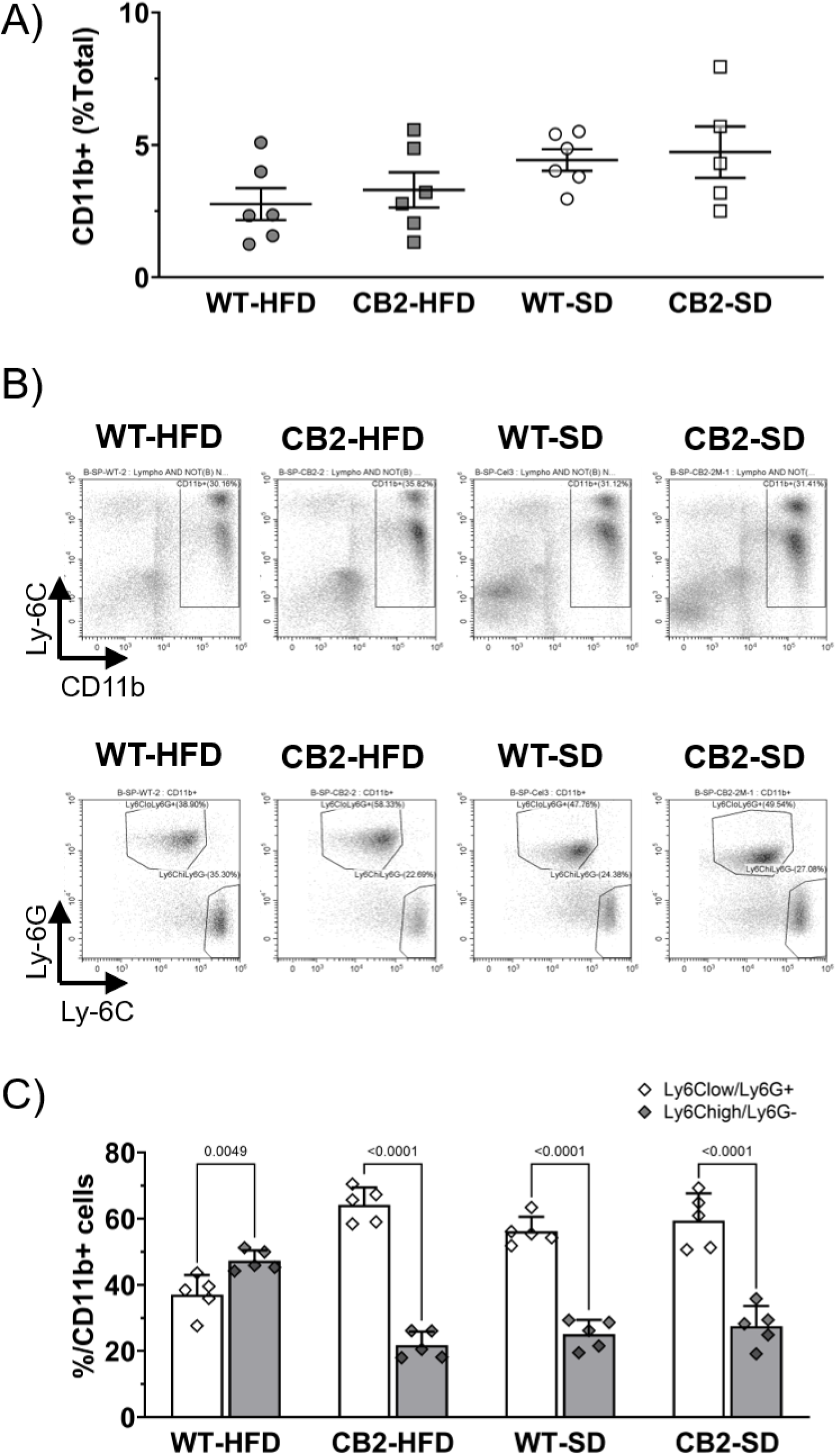
HFD modulates splenic CD11b^+^ myeloid subpopulation in WT animals but not in CB2 knockouts. Population of various immune cells has been analyzed by flow cytometry using spleen. A) Total number of CD5^-^CD19^-^NK1.1^-^CD11b^+^ cells. B) Actual flow cytometry plot of analyzed splenocytes. Top panels shows CD5^-^CD19^-^NK1.1^-^ population gated by CD11b; Bottom panels shows this CD11b^+^ population gated by Ly-6C and Ly-6G. C) Quantification of CD11b^+^ subsets (Ly6C^low^Ly6G^-^ cells and Ly6C^high^Ly6G^+^ cells). Although CD11b^+^ population has unchanged by the genotype or feeding (A), Ly6C^low^Ly6G^+^ subsets as well as Ly6C^high^Ly6G^-^ subsets have been largely modified by HFD in WT animals (B and C). Quantified data are expressed as means ± SEM of 5-6 mice/group. A significant genotype/feeding effect on each cell subpopulation is indicated by one-way ANOVA (A) or three-way ANOVA (C) followed by Tukey’s multiple comparisons test, with respective p-value in the figure.

### CB2 deficiency did not modify CD11b^+^ myeloid subpopulation in peripheral blood and bone marrow after the HFD exposure

While we found the clear genotype difference in CD11b^+^ splenocytes after the HFD exposure, no significant genotype effect to the CD11b^+^ population in bone marrow and peripheral bloods has been found (Fig 6A and C). However, HFD clearly modified the subsets of bone marrow myeloid cells in both WT and CB2 knockouts, which decreased Ly6C^low^Ly6G^+^ cells and parallelly increased Ly6C^high^Ly6G^-^ cells (Fig 6B). Interestingly, this modification pattern was comparable to the splenic myeloid subpopulation in HFD-fed WT animals (Fig 5C). These results suggest that chronic exposure of HFD may anyway lead the modification of myeloid subsets in bone marrow regardless of genotypes.

**Figure 6.**
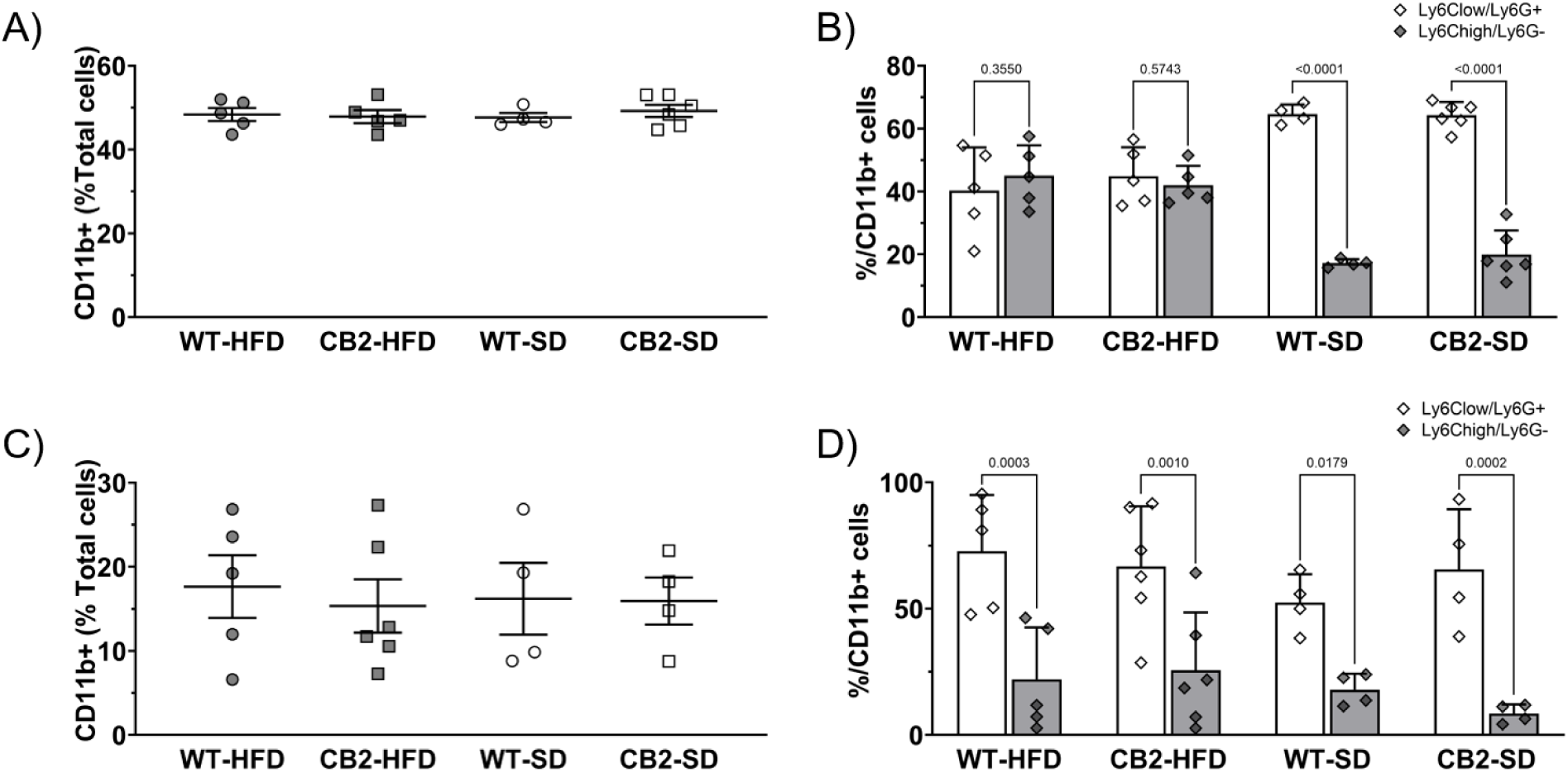
CB2 deficiency did not modify myeloid subpopulation in bone marrow and peripheral blood. Subsets of CD11b^+^ myeloid population has been analyzed by flow cytometry using bone marrow (A, B) or peripheral blood (C, D). 5-weeks HFD exposure significantly modified CD11b^+^ subsets (Ly6C^low^Ly6G^-^ and Ly6C^high^Ly6G^+^) in bone marrow (B), but not in peripheral blood (D). Further, no genotype effect has been observed in CD11b^+^ subsets in bone marrow and peripheral blood. Quantified data are expressed as means ± SEM of 4-6 mice/group. A significant genotype/feeding effect on each cell subpopulation is indicated by one-way ANOVA (A, C) or three-way ANOVA (B, D) followed by Tukey’s multiple comparisons test, with respective p-value in the figure.

We further analyzed same myeloid subsets of peripheral blood, and found that subpopulation of serum myeloid cells has been unchanged by HFD, no matter the CB2 receptors are presence or absence (Fig 6D). This result indicates that myeloid subpopulation modification by HFD has clear priority, that HFD will more affect to myeloid cells of bone marrow, then spleen, and least to them of peripheral blood.

## Discussion

Most reports studying cannabinoid CB2 receptor says that activation of CB2 signaling causes anti-inflammatory effect together which may result as e.g. anti-hyperalgesia in periphery. Such CB2-mediated anti-inflammatory effect has been observed in models of neuropathic and inflammatory pain[15], rheumatoid arthritis[16], multiple schlerosis[17], Duchenne Muscular Dystrophy[18] and even more[19]. Our previous study using CB2 knockout animals showed that lack of CB2 signaling exacerbates neuroinflammation[5–7], which supports anti-inflammatory role of CB2 receptors. However in the present study, we found an exceptional case that CB2 activity may rather be proinflammatory. Thus, peripheral neuroinflammation as well as modification of immune cell population induced by HFD has been significantly diminished in CB2 deficient animals.

Peripheral neuroinflammation has been assessed by 1) pain behavior, 2) macrophage infiltration to peripheral nerve tissue and 3) CXCR4 expression on peripheral nerve tissue. As previously reported, CB2 deficiency has enhanced pain and macrophage infiltration under pSNL-induced nerve injury. We further found the overexpression of C-X-C chemokine receptor CXCR4 in nerve-injured CB2 knockouts. CXCR4 is the most widely expressed chemokine receptors known by its critical role to direct the migration of leukocytes to the site of inflammation [20]. Recent study showed that CXCR4 also takes major role on tissue repair and regeneration in various organs including peripheral nerves[21]. Thus, CXCR4 re-express on the damaged axons among the nerve injury to possibly promote the nerve regeneration[22]. In our study, we have observed the limited expression of CXCR4 around damaged region in nerve-ligated WT animals, whereas CXCR4 positive signal has widely spread out to the non-damaged region in nerve-ligated CB2 knockouts. Overall, our present finding on CXCR4 expression together with hyperalgesia and macrophage infiltration supports that CB2 receptors have anti-inflammatory effect under the nerve-injured neuropathic pain state.

However, all of these three outcomes of enhanced inflammatory response has appeared in WT, but not in CB2 knockouts, after the HFD exposure. 5-week HFD exposure lead significant hyperalgesia, upregulation of macrophage infiltration in nerve tissue and overexpressed CXCR4 in nerve and DRG of WT, whereas none of them were seen in CB2 deficient animals. Our previous study showed that CB2 knockouts reduces the macrophage infiltration into the adipose tissue and inflammatory cytokine levels of liver[23]. Thus CB2 deficiency resists the inflammatory effect of HFD, however its molecular mechanisms still remained unknown.

Our present results added the new insight about 1) effect of CB2 deficiency to the peripheral neuroinflammation and pain, 2) possible contribution of CXCR4 and 3) immune cell profiles including CD11b^+^ myeloid subsets after the sub-chronic HFD exposure. We focused on CXCR4 since past report showed that agonist-bound CXCR4 will form heterodimer with agonist-bound CB2 receptors[24]. CB2-CXCR4 heterodimer has unique role to block Gα13/RhoA-signaling and inhibit the cell migration[25]. HFD itself is known to upregulate the Gα13/RhoA-signaling, to lead various physiological aspects including inflammation, cell transformation, stress fiber formation and myofiber-type remodeling in diet-induced obesity [26], suggesting that Gα13/RhoA-signaling may be one of the key factor that connects HFD, inflammation, CXCR4 and CB2 receptors. However most of the past results shows that inhibition of Gα13/RhoA-signaling, which could be lead by dimerization of CXCR4 and CB2 receptors, is important to resist the HFD effect. As present study showed the contradictory outcome, deeper study about contribution of Gα13/RhoA-signaling to CB2-mediated inflammatory response after HFD exposure may required for the further discussion. Also the expression pattern of CXCR4 in other tissue such as liver and muscle may be important to understand whether CXCR4-mediated Gα13/RhoA-signaling is really the key factor for the protective effect of CB2 deficiency to HFD-evoked systemic inflammation (including neuroinflammation as we have observed in the present study).

Our study also showed that sub-chronic 5-week exposure to HFD will modify the pattern of CD11b^+^ myeloid subsets in spleen and bone marrow. Such HFD-induced myeloid subset modification was affected by CB2 deficiency specifically in spleen, that Ly6C^low^Ly6G^+^ cells has been decreased while another subset Ly6C^high^Ly6G^-^ cells has been increased only in HFD-fed WT animals, and not in HFD-fed CB2 knockouts. Interestingly this HFD-evoked myeloid subset modification happened regardless of genotype in bone marrow. Also note that such myeloid subset modification was not observed in peripheral blood, therefore HFD had no effect to the myeloid profile in the blood. Collectively, these results show that 1) CD11b^+^ cells in bone marrow were affected by HFD regardless of CB2 receptor existence, 2) CD11b^+^ cells in the spleen are affected by HFD due to the presence of CB2 receptors, while 3) CD11b^+^ cells in peripheral blood were not affected by HFD exposure with or without CB2 receptors. These results seem to be inconsistent with the known hematopoietic mobilization circuit [27], starting from bone marrow to the bloodstream then reach to the peripheral organs, therefore at least CD11b^+^ cells in peripheral blood should show comparable subsets as either bone marrow or spleen.

There are couple of possible molecular mechanisms which explain this controversy we have seen in our study. As macrophage infiltration (Fig 2) and inflammation-related CXCR4 expression were modified by CB2 deficiency (Fig 3 and 4), we propose that CB2 signaling may regulate cell infiltration via CXCR4 activity. It is known that adipose tissues become hypertrophied during obesity[28], which may enhance the infiltration of immune cells such as macrophages to the adipose tissues[29]. As CD11b^+^Ly6G^-^Ly6C^high^ cells are mostly monocytic cells which have higher mobility, we propose that CD11b^+^Ly6G^-^Ly6C^high^ cells infiltrated from blood vessel to the surrounding adipose tissues then to the spleen (Fig. 5). Since CXCR4 expression was reduced in HFD-fed CB2 knockouts, mobility of CD11b^+^Ly6G^-^Ly6C^high^ cells may also be reduced, leading to less infiltration and further inhibition of splenic CD11b^+^ subset upregulation.

There are several candidates for two CD11b^+^ subpopulations we found in the present study. Most possible candidate we propose is two subsets of myeloid-derived suppressor cells (MDSC)[30–32]. MDSC consists of two major subsets of Ly6C^low^Ly6G^+^ polymorphonuclear cells (PMN-MDSC) and Ly6C^high^Ly6G^-^ monocytic cells (Mo-MDSC), which is comparable with particular subsets we found in the present study. Since MDSCs are immature myeloid cells, it is known to differentiate into either M1 proinflammatory macrophage and M2 anti-inflammatory macrophage, depending on its pathological conditions[33–35]. In our results, HFD increased Ly6C^high^Ly6G^-^ subsets and decreased Ly6C^low^Ly6G^+^ subsets in the spleen of WT animals, which we observed clear enhancement of pain and neuroinflammation development. Taken together, it is possible that CB2 activity enhanced HFD-evoked inflammatory response in WT animals by modification of MDSC subsets, which may have led upregulation of proinflammatory macrophages and downregulation of anti-inflammatory macrophages. However this is still a speculation and requires detailed analysis of two CD11b^+^ myeloid subpopulations found in this study, together with deeper evaluation of relating immune cells in wide range of immune-related organs including blood, bone marrow and adipose tissues.

## CONCLUSION

In summary, this paper reports not the anti-inflammatory, but rather the pro-inflammatory role of CB2 receptors which is observed after 5-week sub-chronic HFD exposure. While pain and peripheral neuroinflammation has been exacerbated in nerve-injured CB2 deficient animals, HFD evoked pain and neuroinflammation in WT, not in CB2 knockouts. Interestingly, HFD modified splenic CD11b^+^ monocyte subsets are modified only in WT animals; Ly6C^high^Ly6G^-^ subsets has been increased whereas Ly6C^low^Ly6G^+^ subsets decreased. Overall results suggest that HFD leads peripheral neuroinflammation through CB2 signaling, possibly by modification of CD11b^+^ myeloids that may contribute to the inflammatory response. Further, this study indicates CB2 activity is not always anti-inflammatory and may have totally different role on systemic inflammation development.

## Competing interests

All authors declare that they have no competing interests around the present research.

## Authors’ contributions

C.N. designed the experiments. C.N. and H.H. performed the experiments and the data analysis. C.N., H.H. and T.A. drafted the manuscript. All authors contributed to and have approved the final manuscript.

## Acknowledgment

This research was supported by the Deutsche Forschungsgemeinschaft (DFG Eigene Stelle 1198/2-1) and Japan society for promotion of science (JSPS Home-Returning Researcher Development Research 19K24693). We also thank Andreas Zimmer for providing us the animals. We further appreciate technical assistance from Astrid Markert, Kerstin Michel, Kaede Ito, Yuya Kasai and Hsiao-Hsieh Wang.

## Notes

### Competing Interest Statement

The authors have declared no competing interest.

